# Laboratory yeast crosses reveal limited epistasis in the genetic basis of complex traits

**DOI:** 10.64898/2026.04.04.716439

**Authors:** Misha Gupta, Caroline M Holmes, Jenya Belousova, Shreyas Gopalakrishnan, Artur Rego-Costa, Michael M Desai

## Abstract

Mapping the genetic basis of complex traits is complicated by the presence of epistatic interactions between loci. While work in molecular genetics identifies numerous specific genetic interactions, statistical analyses of quantitative traits frequently conclude that additive (nonepistatic) models explain most heritable variation. However, these conclusions are typically limited by the narrow range of genetic relatedness(e.g. in F1 offspring of a biparental or circular cross). Here, we use a barcoded panel of *Saccharomyces cerevisiae* genotypes with a broad range of relatedness to quantify the effects of epistasis on the genetic architecture of seven complex traits. We find limited contributions of epistasis to the genetic basis of these traits. These results indicate that epistasis beyond that detected in standard yeast crosses may exist, yet it contributes little to phenotypic variance in these systems.

## Introduction

Mapping the genetic basis of complex traits is a central goal in genetics. Epistasis, in which loci interact to affect phenotypes, can complicate efforts to infer these genotype-phenotype maps. Numerous studies in model organisms like budding yeast have used quantitative-trait-locus mapping to quantify contributions of additive and epistatic terms to phenotypes[1,2]. These studies have consistently found that the genetic basis of complex traits is dominated by additive terms, with epistasis typically playing a detectable but weak role[3,4,5]. This body of work stands in contrast to molecular studies that directly measure genetic interactions (e.g. double-deletion studies in yeast[6,7]), which find strong signatures of epistasis.

However, quantitative genetics studies in model organisms usually involve individuals with narrow relatedness structures. Numerous studies have focused on F1 offspring of a biparental yeast cross[1]; these offspring all share half of their genomes with each other. Other studies analyzed deeper crosses or those involving more parents[8], or recombinant inbred lines[9]. The resulting offspring still span a range of relatedness values that may differ between studies, yet is individually narrow. This limits their power to use variance partitioning or related methods to estimate the contribution of epistasis to phenotypic variance.

Here, we construct a panel of barcoded *Saccharomyces cerevisiae* individuals generated from F1 segregants of a biparental cross[10] and two backcrosses of similar segregants (one to each original parent). We measure seven relative growth rate phenotypes for each segregant, and quantify the effects of epistasis on the genetic architecture of each trait. The broader relatedness values between these segregants should allow us to detect signatures of epistasis beyond those observed from the F1 segregants alone. However, we find that these effects are weak and explain only a tiny fraction of phenotypic variation.

## Results

We selected 1480 barcoded haploid F1 offspring from a previous cross[10] between strains derived from a laboratory budding yeast strain, BY4741, and a wine strain, RM11-1a[10]. For the ∼100,000 F1 offspring from the cross, fitness was measured in multiple environments. We chose two sets of 732 segregants from those sequenced at high coverage. We then backcrossed half of the first set with the BY parent, and half with the RM parent. This generated a combined panel consisting of the second set of 732 F1 segregants (F1-R), 332 BY-backcrossed segregants (BY-BX), and 332 RM-backcrossed segregants (RM-BX), each with a unique DNA barcode. We sequenced each segregant and inferred complete genotypes using a Hidden Markov Model[10], exploiting the recombination structure of the segregants. We measured the relative growth rates of each segregant in seven different media conditions by tracking changes in barcode frequencies over time(Fig.1; see Methods), yielding measured fitness values of RM-BX and BY-BX as well as re-measured fitness values of F1-R.

We compared the measured phenotypes of our segregants to the predictions from a model of linear effects and pairwise interactions inferred from the full set of the previous 100,000 F1 segregants[10]. If epistasis beyond that captured in the F1 panel was widespread, predictions derived from F1 data should perform poorly in backcrossed segregants, where allele frequencies and genetic backgrounds differ.

In most environments, we see differences in model performance between segregants that were in the training data (F1-R), and segregants of BY-BX and RM-BX that were not previously seen by the model (Fig 2, leftmost column). Normalised Root Mean Squared Error (RMSE) is generally higher for RM-BX segregants, which occupy higher-fitness regimes. However, Fig 2 (center column) shows that when conditioning on fitness, predictive accuracy is similar across genetic backgrounds. The class-specific differences arise from the distribution of phenotypic values rather than genotypic class itself. This pattern is consistent with diminishing returns epistasis, where mutational effects are compressed at higher fitness. In contrast, in GuCl and YNB, differences persist across genotype classes even after controlling for fitness (Fig 2, center column). The model trained on just F1 segregants fails to capture class-dependent deviations present in these populations. Prediction error cannot be explained only by phenotype value; these deviations suggest higher-order epistatic interactions that are not fully represented in the training data.

**Figure 1.**
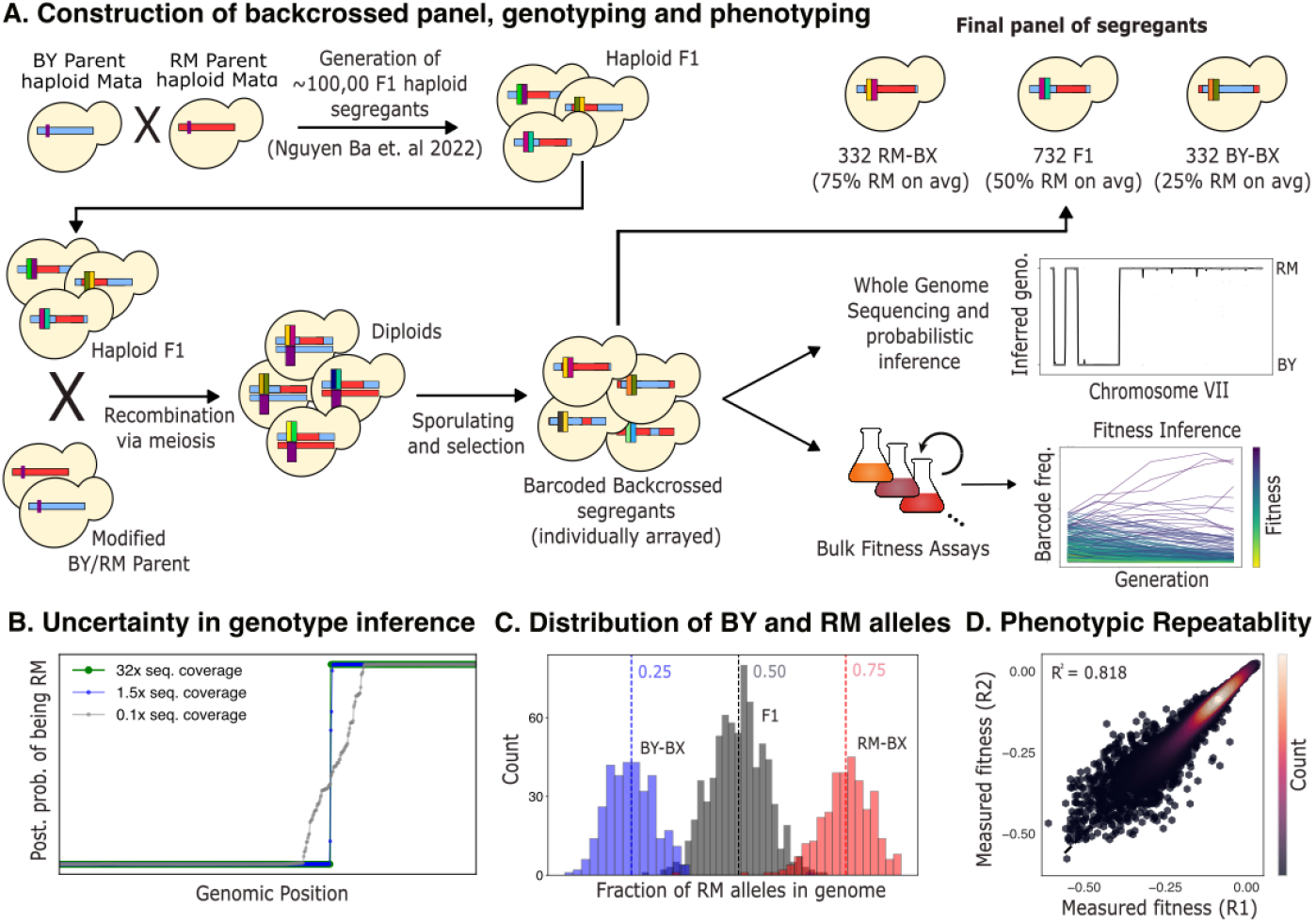
We built a panel of barcoded segregants from an F1 cross and by back-crossing these segregants to their parents. **A.** Construction of the segregant panel involved mating barcoded haploid F1 segregants with either the BY or RM parent, sporulating the diploids and selecting for haploids with the barcode maintained. **B**. Comparison of putatively inferred genome of a single segregant (sequenced at 35x coverage), the inferred genome from data downsampled to 1.5x coverage (minimum threshold for panel) and 0.1x for comparison. Full panel errors in genotyping can be seen in Fig. S1. **C**. Fraction of RM alleles in each of the two backcrosses and the original F1 cross. **D**. Repeatability of phenotypic measurements in all environments combined. See Fig. S2 for analogous plots for each environment individually.

**Figure 2.**
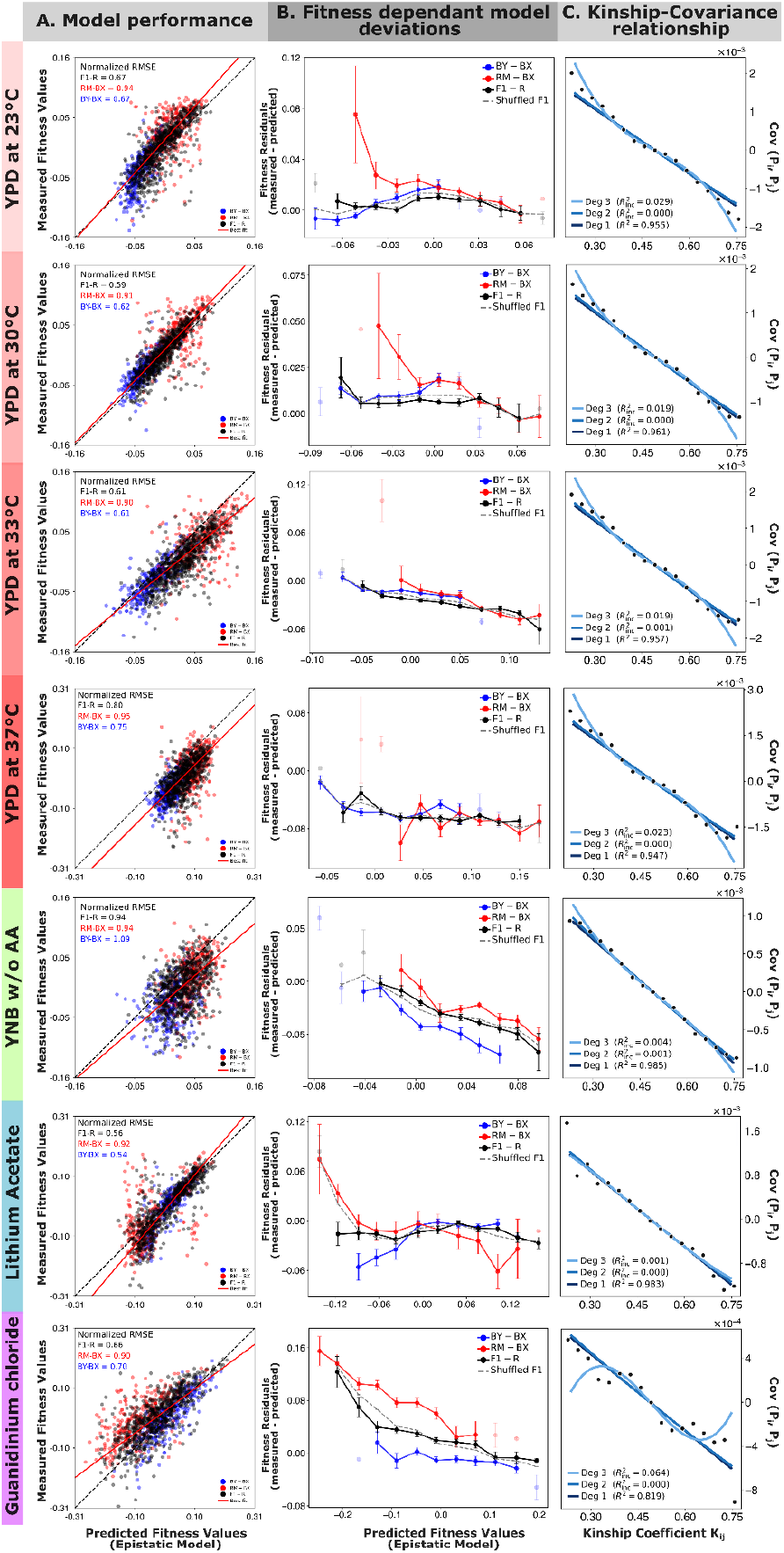
**A.** Predicted and experimentally measured fitness show slight systematic deviations from the 1:1 line (dashed black). The density-weighted best-fit line (red) differs marginally in slope; the offset reflects an inference artifact and differences in mean population fitness. Within each condition, the segregant with median fitness was set to zero, with all other values shifted accordingly. Normalized RMSE to the best-fit line is reported. **B**. Residuals between measured fitness values and fitness values predicted by the epistatic model in a particular fitness bin (calculated as the mean of the bin). If a given bin has fewer than 5 points, it is faded out. The gray dashed line shows the null distribution, computed by shuffling strain labels within predicted-fitness bins over 1,000 iterations (F1-R shown; shuffled BY-BX and RM-BX are equivalent). GuCl and YNB show systematic differences in model performance between the different genetic backgrounds. Since each bin has been sliced by predicted fitness value, this accounts for the systematic differences in the fitness values of BY-BX and RM-BX. **C**. The linear model explains the majority of variance in phenotypic covariance as a function of a kinship coefficient, across all conditions with higher-order terms contributing minimally, as reported by the incremental variance explained 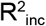 Any bin with less than 100 pairs of points was excluded from plotting and fitting.

To avoid relying on the assumptions of any particular model, we examine the relationship between kinship and phenotypic covariance; nonlinearity in this relationship is a signature of epistasis[11,12,13]. The linear model fits the data well across all conditions (R^2^>0.94 in all conditions except GuCl), and higher-order polynomial terms explain little additional variance 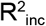, consistent with an predominantly additive genetic architecture.

## Discussion

We built and phenotyped a lab cross spanning genetic relatednesses to test whether the apparent lack of strong epistasis in previous QTL studies was due to the limited relatedness studied. We found minimal evidence for additional epistasis in this new cross. When higher-order epistasis is present in our laboratory environments, its effect is modest, and does not seem to drive trait prediction. In the specific context of two growth environments, we were able to detect deviations expected from higher-order epistasis that the predictive model did not include.

Even when the structure of our panel is designed to detect non-additive variance, the signal is absent in the majority of our growth environments, at least on the scale of our lab experiments. Instead, linearity indicates that higher-order terms contribute little to measurably affect the expected additive relationship[14]. It seems that additive or low-order epistatic models are likely sufficient for the majority of phenotypic prediction tasks in yeast laboratory crosses, at least for the growth assays. However, higher-order epistasis can be detected and may matter in specific contexts. This may help explain why predictive accuracy in studies often plateaus despite large sample sizes. The remaining phenotypic variance seems to not be driven by strong higher-order epistatic terms, but rather by a combination of weak interactions, measurement noise, and other as yet unmodeled environmental or technical factors.

## Materials and Methods

We selected 1480 barcoded haploid F1 offspring from a cross between strains derived from BY4741 and RM11-1a, both of which were converted to Mat**α**. These F1 individuals each contained a barcode at the HO locus. Of these, 372 were backcrossed to the BY4741 parent and 376 were crossed with the RM11-1a parent; the diploids were selected, sporulated and one Mata clone was retained from each new cross. These offspring maintain the barcode of their F1 parent, and therefore do not need to be rebarcoded. We then performed whole-genome sequencing on these strains. Large, continuous blocks of the genome are inherited from either the RM or BY parent, which allows us to infer the entire genome even from ‘low’ coverage sequencing. As in[10], we inferred the genotype of any unsequenced regions using a Hidden Markov Model, removing any segregants with <1.5x coverage or signatures of diploidy. Our final panel then consisted of 664 backcrossed individuals and an additional 732 F1s from our original cross, all with unique barcodes at the HO locus.

We measured phenotypes in bulk fitness assays with two replicates. We did this in seven different growth media: YNB (minimal yeast media), a temperature gradient of 23C, 30C, 33C, and 37C in YPD (rich media), lithium acetate (salt stress), and guanidinium hydrochloride (salt stress). In each growth medium, we passaged 671 μL of saturated culture into 100 mL of fresh media each day, which resulted in a 2^7^-fold daily dilution for 7 days. After days 1, 2, 4, 6, and 7, we took a sample of the cells, PCR amplified the barcoded region, and sequenced the amplicon DNA. We used that data to measure the relative frequencies of each barcode lineage at each timepoint.From these frequency trajectories, we inferred the relative growth rates of each barcoded segregant using a maximum-likelihood model. See SI for further information and extended protocols.

For the kinship-based analysis, we computed the pairwise kinship coefficient K_ij_ from genome-wide Hamming distances, binned pairs into 25 equal-width bins, and calculated the phenotypic covariance within each bin. Weighted polynomial regression (weighted by bin size) was fitted to the binned covariance estimates. We estimate goodness of fit by calculating a weighted R^2^ to the linear model, and estimate incremental variance explained by non-linearities for the higher degree models as 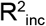

## Data archival and availability

Fitness values and accompanying code to be released upon publication on github (archived at Zenodo). Raw sequencing reads have been deposited at NCBI SRA.

## Acknowledgments

The authors would like to thank Joao Ascensao for his help and comments, grant R01-GM104239 from the NIH for funding, and members of the Harvard Bauer Core facilities for help with sequencing. CMH was funded by the Jane Coffin Childs Memorial Fellowship. The computations in this paper were run on the FASRC Cannon cluster supported by the FAS Division of Science Research Computing Group at Harvard University.

## References

1. Bloom, Joshua S., et al. “Finding the sources of missing heritability in a yeast cross.”Nature 494.7436 (2013): 234–237.

2. Ehrenreich, Ian M., et al. “Dissection of genetically complex traits with extremely large pools of yeast segregants.”Nature 464.7291 (2010): 1039–1042.

3. Bloom, Joshua S., et al. “Genetic interactions contribute less than additive effects to quantitative trait variation in yeast.”Nature communications 6.1 (2015): 8712.

4. Buzby, Cassandra, et al. “Epistasis and cryptic QTL identified using modified bulk segregant analysis of copper resistance in budding yeast.”Genetics 229.4 (2025): iyaf026

5. Matsui, Takeshi, et al. “The interplay of additivity, dominance, and epistasis on fitness in a diploid yeast cross.”Nature Communications 13.1 (2022): 1463.

6. Costanzo, Michael, et al. “A global genetic interaction network maps a wiring diagram of cellular function.”Science 353.6306 (2016): aaf1420.

7. Shen, John Paul, et al. “Combinatorial CRISPR–Cas9 screens for de novo mapping of genetic interactions.”Nature methods 14.6 (2017): 573–576.

8. Cubillos, Francisco A., et al. “High-resolution mapping of complex traits with a four-parent advanced intercross yeast population.”Genetics 195.3 (2013): 1141–1155.

9. Broman, K. W. “The genomes of recombinant inbred lines.”Genetics 173.4 (2006): 2419–2419.

10. Nguyen Ba, Alex N., et al. “Barcoded bulk QTL mapping reveals highly polygenic and epistatic architecture of complex traits in yeast.”Elife 11 (2022): e73983.

11. Fisher, Ronald A. “XV.—The correlation between relatives on the supposition of Mendelian inheritance.”Earth and Environmental Science Transactions of the Royal Society of Edinburgh 52.2 (1919): 399–433

12. Wright, Sewall. “Coefficients of inbreeding and relationship.”The American Naturalist 56.645 (1922): 330–338.

13. Young, Alexander I., and Richard Durbin. “Estimation of epistatic variance components and heritability in founder populations and crosses.”Genetics 198.4 (2014): 1405–1416.

14. Falconer DS, Mackay TFC (1996) Introduction to quantitative genetics, 4th Ed. (Longman, Essex, UK)

