## Supplemental Information for "Laboratory yeast crosses reveal limited epistasis in the genetic basis of complex traits"

### Supporting Information Text

**Construction of backcross panel.** All the F1 parental strains were generated in a previous study (1). The BY and RM parental strains were derived from BY4741 $S288C : MATa, his3\Delta1, ura3\Delta0, leu2\Delta0, met17\Delta0$ and RM11-1a (a haploid derivative of Robert Mortimer's Bb32(3): $MATa, ura3\Delta0, leu2\Delta0, ho :: KanMX$) (2), respectively. These strains were further modified for a barcoding scheme (1) to have the complete barcode and URA3 gene at the HO locus (MATa, $ura3\Delta0$, $leu2\Delta0, his3\Delta1, can1 :: Ste2pr\_SpHIS5, HO :: LEU2 :: 1/2URA3\_bc1 :: bc2\_2/2URA3$). We further modified these BY and RM parents for our experiment, allowing us to select against unmated F1s, remove the BY/RM barcode, and maintain the F1 barcode in the final panel.

First, we replaced the HO locus in BY and RM with a HphMX cassette, amplified with overhangs that are homologous to the HO locus from strains in a different study (3). A modified version of the Gietz and Schiestl protocol (4) was used to transform the linear HphMX cassette into 5 ml of cells that were grown to $OD_{600} = 0.8$ and then pelleted (BY and RM separately). The cell pellet was washed with sterile, molecular grade water three times, and mixed with the transformation mix (240 $\mu$L of PEG3350 (50% (w/v)), 36 $\mu$L of 1M Lithium Acetate, 10 $\mu$L of 10 mg/ml single stranded carrier DNA (Sheared Salmon Sperm - Invitrogen), 1 $ug$ of our linear fragment and sterile, molecular grade water to top up volume to 360 $\mu$L). The transformation reaction was incubated at 42C for 30 minutes, allowed to grow in 1 ml YPD (1% Difco yeast extract, 2% Gibco Bacto peptone, 2% BD Difco dextrose) for 1 hour to express the resistance cassette, and then plated on selective media (YPD + 2% agar + 300$\mu$g/ml of HygromycinB) overnight. Cells were streaked to single colonies to recover our BY*/RM* parental genotypes ($MATa, ura3\Delta0, leu2\Delta0, his3\Delta1, can1 :: Ste2pr\_SpHIS5, HO :: HphMX$). The mating type switch for BY*/RM* parents was performed using a plasmid containing the HO endonuclease, using the protocol and plasmid $pAN216a - GAL :: HO - Ste2pr :: SkHIS3 - Ste3pr :: LEU2$ from (1) and galactose induction of the mating switch. The final parental genotypes, BYref/RMref $MAT\alpha, ura3\Delta0, leu2\Delta0, his3\Delta1, can1 :: Ste2pr\_SpHIS5, HO :: HphMX$ are no longer uracil auxotrophs, contain no barcodes and can mate with the MATa F1 segregants.

Each F1 segregant and BYref/RMref was separately grown to saturation. For each mating, 100 $\mu$L of the F1 parent as well as the chosen BYref/RMref parent was added to a 96-well, deep-well plate, and pelleted down. Each pellet was resuspended in 20 $\mu$L of sterile water, and 4 $\mu$L was spotted on an omni-plate. This was incubated at 30C to allow mating to occur, for 48 hours, or till colonies were seen. At this point, each growing colony contains a mixture of unmated BYref/RMref, unmated F1s (sensitive to HygromycinB) and the desired diploids that have undergone recombination. Each colony was grown in 400 $\mu$L of YPD + HygromycinB (300$\mu g/ml$) in deep well plates to remove any unmated F1s from the mixture. We used mild sonication to recover haploids by diluting 20 $\mu$L of the fully grown culture in 480 $\mu$L pre-sporulation media (1% yeast extract, 2% Bacto peptone, 1% potassium acetate, 0.005% zinc acetate), growing for 24 hours at room temperature, in deep well plates with AeraSeal breathable films (Sigma Aldrich), pelleting and resuspending into 500 $\mu$L of sporulation media (1% potassium acetate, 0.005% zinc acetate), and growing for 72 hours at room temperature. The mixture contains unmated F1s that are uracil auxotrophs as well as prototrophic backcrossed segregants. The spores were germinated by pelleting 200 $\mu$L spores and digesting their asci by resuspending in 50 $\mu$L of 5mg/mL Zymolyase 20T (Nacalai Tesque) containing 20 mM DTT (Thermo Fisher) for 15 min at 37C. The tetrads were disrupted by sonication (3 cycles of 30 seconds at 80% power, with 30 seconds on ice between each cycle) and the cells were recovered in media lacking uracil, histidine and leucine (to remove any diploids that may have leaked through) (6.71 g/L BD Difco Yeast Nitrogen Base, complete amino acid supplements (Thermo Fisher) lacking uracil, histidine, leucine, 2% BD Difco dextrose). These are then streaked to single colonies on the same selective media + 2% agar plates, allowing for 3-4 days of growth. We repeat this process for 384 F1 strains with BYref and different 384 F1 strains with RMref, individually array each clone, and store at -80C in 15% glycerol as a cryoprotectant.

**Whole genome sequencing and genome inference.** Using previous methods for genome imputation and knowledge of breakpoints (1) in our segregants, we can get clean genomic data with 'low' sequencing coverage. We used modified DNA extraction protocols adapted from (5) and sequencing protocols modified from (6). Each individual segregant from the panel was grown in 1 ml of YPD as liquid culture to saturation, and 300 $\mu$L of culture was pelleted in deep well plates. The DNA extraction was performed on Beckman Coulter Biomek FxP robots. Each cell pellet was resuspended in 50 $\mu$L of zymolyase buffer (5 mg/mL Zymolyase 20T (Nacalai Tesque), 1 M Sorbitol, 100 mM Sodium Phosphate pH 7.4, 10 mM EDTA, 0.5% 3-(N,N-Dimethylmyristylammonio)-propanesulfonate (Sigma, T7763), 200 $\mu$g/mL RNase A, and 20 mM DTT(Thermo Fisher)), thoroughly vortexed and incubated at 37C for 45 minutes. 85 $\mu$L of the modified 'BOMB' buffer from (5) (4M guanidinium-isothiocyanate (Goldbio G-210–500), 50 mM Tris-HCl pH 8, 20 mM EDTA) and 120 $\mu$L of isopropanol (VWR BDH1133-4LP) was added. This was mixed for 3 minutes after each step. 20 $\mu$L of Zymo Research MagBinding beads were added to bind the DNA and mixed for 3 minutes. The beads were separated from the debris using a Magnum FLX 96-well magnetic separation rack (Alpaqua), and the supernatant removed. The beads were washed with 400 $\mu$L of isopropanol and twice with 300 $\mu$L of 80% ethanol to remove cell debris, and left open for 10 minutes for alcohol to evaporate. We then added 75 $\mu$L of sterile, molecular grade water to beads to get our DNA into solution, separated the beads using a same magnetic rack, and 44 $\mu$L of the supernatant (containing the DNA) was added to 96-well PCR plates (Bio-Rad HSP9631) for library preparation and storage. All segregrants were treated separately.

For efficient downstream sequencing, we used the protocol elucidated in (6). Each sample was quantified using the AccuGreen Broad Range dsDNA Quantitation Kit (Biotium) using the Spectramax i3 plate reader (509 nm/530 nm), and all samples were normalized to a DNA concentration of 2 ng/$\mu$L using the Mantis (Formulatrix) for small volume liquid handling. Once at the appropriate concentration, the extracted DNA was sheared and amplified using the Illumina tagmentase (TDE1 enzyme,

Illumina 20034198) and exact protocol from (6). Since each DNA sample was run with a unique combination of Illumina Nextera adapters, we could pool all samples into a single pool and de-multiplex at a later stage. Fragments were pooled by equal weight into appropriate pool. Double sided size selection was performed to clean up the reaction and get fragments of around 300 bp length, for optimal sequencing (0.55-0.85x cleanups). The prepped libraries were sequenced (paired end) on a single lane of NovaSeq SP. We sequenced each individual to low coverage, and then use probabilistic imputation to get the full genomes. The location and identity of all SNPs in our segregants is known from deep sequencing of the parental BY/RM (1) where after curation, the two parents differ at only 41,594 SNPs. The final reference genomes and SNP positions can be found in (1) and at the accompanying GitHub repository. From here, we only need to infer recombination breakpoints to get the entire genome for each of our segregants (7).

For each sample, we use Trimmomatic (version 0.39) to clean raw reads from a FASTQ file, use bowtie2 (8) (version 2.5.1) to align our trimmed reads to the 2 different yeast genomes, and then count how many reads support a given allele (BY/RM) at a given SNP. We impute SNP genotypes from these counts using the exact Hidden Markov Model from (1), using information about recombination and sequencing error rates. We enforce atleast 1.5x coverage to ensure sufficient data for correct imputations. For each individual, we thus produce a 'probabilistic genotype', using the convention that 1 indicates the RM allele and 0 the BY allele. For the error rates in the model, we use probabilities informed by knowledge of sequencing errors and average recombination rates across the genome. All values are hence taken from (1) since the sequencing chemistry and organism is the same. Our genomes have low uncertainty, even at low coverage levels. We check this by downsampling our data and comparing inferred genotypes from low (1.5x) and high (full) sequencing coverage of each clone. The errors per SNP in posterior probability (calculated as the difference between high and low-coverage posterior probabilities) is $< 0.1$ across all coverage values, as seen in S1. Note that our analysis does not include any genotypes that had coverage <1.5x, were found to be diploids (supporting both the RM and BY allele at a given locus with probability $\sim 0.5$), or had fuzzy recombination breakpoints.

**Bulk fitness assays and amplicon sequencing.** We chose a complex 'fitness' as our phenotype, measuring it in 7 growth environments Table S1. We can track each lineage with its identifying barcode and infer relative fitness effects from these changes in barcode frequencies (9). While this is not exactly the same as direct growth rate measurements in log phase, our measurement tracks frequency changes, and infers fitness relative to the population. We conduct pooled, all-to-all competition assays, by combining all lineages in a single flask per environment, passaging them over time, measuring changing frequencies by counting identifying barcodes, and deeply sequencing to get correct estimates of frequencies. Since each barcode is associated with a known genotype, our inferred fitness values can be correctly assigned.

| Name | Description |
|---|---|
| 23C | YPD at 23C |
| YPD | YPD at 30C |
| 33C | YPD at 33C |
| 37C | YPD at 37C |
| YNB | YNB w/o AA and w/o ammonium sulphate at 30C |
| LiAc | YPD + 20 mM LiAc at 30C |
| GuCl | YPD + 6 mM GuCl at 30C |

**Table S1. Details of different media used as environments, all contain 100$\mu$g/ml Ampicillin**

For each plate of F1s, we combine equal volumes of each segregant from a plate and store in 15% glycerol. Each pool was grown in YPD at $2^5$ dilution to help recover from physiological shocks of a freeze-thaw cycle; all pools are then combined and used to inoculate our fitness assays, each in the appropriate growth media. We inoculate 100 ml of each growth medium with 781 $\mu$L of saturated culture in a 500 ml baffled flask. These passages (781 $\mu$L into 100 ml of media) are performed every 24 hours, resulting in approximately 7 generations of growth per passage (3). We passage for 49 generations, collecting 3 ml of culture each time to sample barcodes. We only extract DNA and sequence at generations 7, 14, 28, 42 and 49.

To extract DNA from a 1.5 ml frozen pellet, we use a silica-column based method (3). Cell pellets were thawed and lysed in 150$\mu$L of a zymolyase lysis buffer (5 mg/mL Zymolyase 20T (Nacalai Tesque), 1 M sorbitol, 100 mM sodium phosphate pH 7.4, 10 mM EDTA, 0.5% 3-(N,N-Dimethylmyristylammonio)propanesulfonate (Sigma, T7763), 200 µg/mL RNAse A, and 20 mM DTT) at 37C for 1 hour. The lysed cells were bound to silica columns (IBI Scientific Mini Genomic DNA Columns IB47207) with 400$\mu$L of a binding buffer (100 mM MES pH 5, 4.125 M guanidine thiocyanate, 25% isopropanol, 6 mM NaOH and 10 mM EDTA) and adding to the column. The column is washed with 400 $\mu$L of an alcoholic buffer (10% guanidine thiocyanate, 25% isopropanol, and 10 mM EDTA), then washed with 600$\mu$L of a wash buffer (80% ethanol, 10 mM Tris pH 8) dried by spinning down at 16000 RPM for 3 minutes, and DNA eluted in 50$\mu$L of sterile water. Each sample was sequenced by tagging with a unique combination of Illumina Nextera adapters and pooling. Our two PCRs to amplify the barcode region allows us to attach unique molecular indices (UMI) to each sample and correct for any PCR bias. For PCR1, dual indexed primers were used that attach at the KanP1 (forward) and HygP1 (reverse) sites in the artificial construct created for the barcode. Each primer contains 8 base pair random UMIs, and tagging indices of variable length.

The PCR was performed with a high fidelity Q5 polymerase (10$\mu$L of extracted DNA, 2.5$\mu$L of 4 mM primers each, 4$\mu$L of 5x Q5 buffer, 0.4$\mu$L of 10 mM dNTPs, and 0.2$\mu$L of Q5 enzyme (NEB M0491L), topping up the volume to 20$\mu$L with sterile, molecular grade water). PCR amplification was performed with the following thermal cycling conditions: Initial denaturation at 98C for 1 minute, followed by 2 cycles of: denaturation at 98C for 10 seconds, annealing at 50C for 10 seconds, extension at 72C for 30 seconds and final extension at 72C for 30 seconds. 20$\mu$L of E. coli carrier mRNA was added to the PCR product and cleaned up using Aline beads (left sided, 50$\mu$L of Aline beads for a 1.25x reaction, eluted in 34$\mu$L of sterile water). All of the eluted, cleaned up DNA was used as the input for the second PCR reaction to add Nextera adapters for sequencing. Custom primers (P5mod and P7mod) were used for sequencing. Kapa (Roche Sequencing KK2502) was used for the final PCR (10$\mu$L of 5x Kapa buffer, 1$\mu$L of 10mM dNTP's, 2.5$\mu$L of 10mM forward and reverse primers, 33$\mu$L of template DNA, 1$\mu$L of the polymerase and sterile, molecular water to top up to 50$\mu$L reaction volume). PCR amplification was performed with the following thermal cycling conditions: Initial denaturation at 98C for 30 seconds, followed by 34 cycles of: denaturation at 98C for 10 seconds, annealing at 62C for 10 seconds, extension at 72C for 30 seconds and final extension at 72C for 30 seconds. Post verification on a gel, the samples were cleaned up using a left sided 0.85x Aline bead cleanup, quantified and pooled equimolar. All samples were run on a NextSeq 1000 P1 (v1.5 chemistry), paired end 300 cycle kit, with 6 bp indices and all extra indices being allotted to the first read.

The experimental, sequencing based readout is a count of each lineage present at a given timepoint. We process raw reads from the sequencer using regex-based python scripts, matching known constant regions around the variable parts of the read. The resulting barcodes are error corrected against a known list, duplicated reads are removed, and each barcode with a unique UMI is counted. We infer fitness values using a ML based approach that fits initial frequencies and fitness effects of each barcode to best explain our data. This fitness inference is performed exactly as in Appendix 2 of (1), using a multinomial model with gaussian prior, all data (joint across replicates) and correcting for standard errors by calculating overdispersion. We measure everything in duplicates, keeping separate batches of media per replicate. We find our measurements are robust to experimental noise. Remeasured F1 fitness values are also in agreement with the previous experiment (1). The robustness of our phenotypic measurements can be seen in Figure S2.

**Prediction of fitness values.** The models we used to predict fitness were trained in (1) and can be found in the accompanying datasets. For each growth environment, fitness values were predicted separately. For the epistatic model, the interaction contributions were added on top of the additive weights (adjusted accordingly in the original training). Following these predictions, a random barcode in the bulk of the measured values was chosen and arbitrarily set to 0. The corresponding shift in fitness values was then applied to all measured values. The same barcode was then shifted to 0 in the predicted values, and the corresponding shift applied to all predicted values. This allows us to account for arbitrary offsets in fitness inference, and is an appropriate method since our fitness is inferred relative to a randomly chosen reference anyway.

**Theoretical details of variance partitioning.** Phenotypic variance can be partitioned into its many components (Variance due to additivity $V_A$, variance due to dominance $V_D$, variance to environmental effects $V_E$ and variance due to higher order epistatic terms $V_I$) simply as $V_p = V_A + V_E + V_D + V_I + \dots$ (10). The works of (11) further expand these concepts by defining a kinship coefficient $K_{ij}$ between 2 individuals $i$ and $j$, defined as the probability that a random allele from these individuals is Identical By Descent. This allows us to write down

$$\mathrm{Cov}(A_i, A_j) \propto V_A K_{ij}, \tag{1}$$

where $A_i$ is the additive phenotype, $V_A$ is the additive variance and $K_{ij}$ is the kinship coefficient. The proportionality factor is set by ploidy. To generalize, we can write this as $y_i = A_i + I_i + E_i + D_i$, where $y_i$ is the phenotypic trait value, $A_i$ is the additive trait value, $I_i$ is the contribution of higher order interactions to the phenotypic trait value, and $E_i$ is the contribution of environmental conditions to the trait value. In our haploid batch culture, $D_i \to 0$ and $E_i$ is minimized. If there are any non-zero interaction terms, they will show up as higher order polynomial terms. Any non-linearities in the relationship between the phenotypic covariance and kinship coefficient will indicate variance explained and trait prediction that can be attributed to epistatic terms. We make the assumption that there are no epistatic interactions or linkage disequilibrium, which would ensure that Equation 1 holds true. Any deviations from linearity of that relationship will point towards our assumption being violated, and the presence of higher order epistatic terms.

***Variance partitioning and epistasis detection.*** For each pair $(i, j)$ of barcodes, we have a kinship coefficient $K_{ij}$, calculated as the Hamming distance normalized by the number of SNPs, and phenotypic measurements $y_i$ and $y_j$. We partition the range of $K_{ij}$ into 25 equally spaced bins and exclude any bin containing fewer than $n_{\min} = 100$ pairs. For each retained bin, we compute the mean kinship and the empirical phenotypic covariance.

We then fit weighted least squares (WLS) polynomials of degree $k \in \{1, 2, 3\}$ to the binned data, weighting each bin by number of points. To assess goodness of fit, we compute the weighted $R^2$ to the linear fit, as $R^2 = 1 - \mathrm{RSS}_1/\mathrm{TSS}$. The incremental $R^2$ of a degree-$k$ polynomial over the linear model is calculated as

$$R^2_{\mathrm{inc}}(k) = \frac{\mathrm{RSS}_1 - \mathrm{RSS}_k}{\mathrm{TSS}}, \qquad k \in \{2, 3\}, \tag{2}$$

where $\mathrm{RSS}_k$ is the weighted residual sum of squares of the degree-$k$ fit, and TSS is the weighted total sum of squares. This quantifies the additional variance in phenotypic covariance explained by the nonlinear terms.

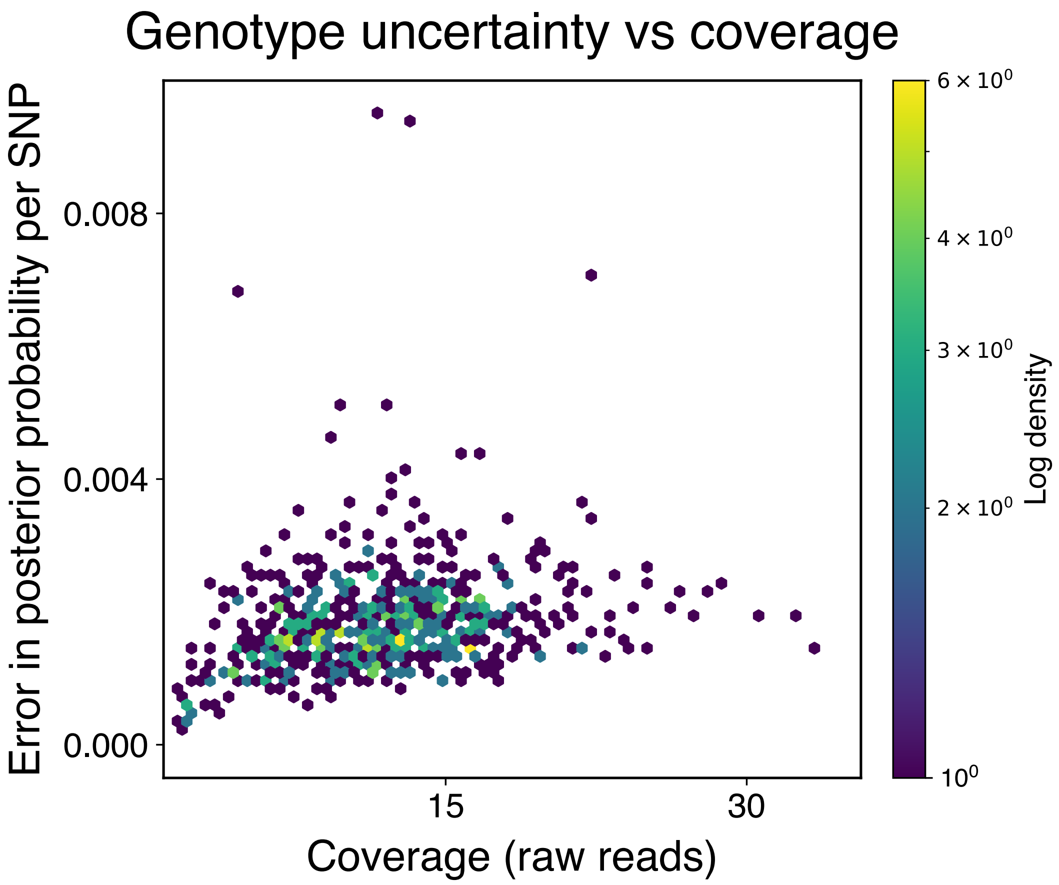


**Fig. S1.** Genome uncertainty as a function of sequencing coverage. The error has been calculated by comparing to data down-sampled to 1.5x coverage, and taking the difference in calculated posterior probabilities. There are 26 outliers that are outside the vertical bounds that have been excluded for visualization. We do not see a difference in genomes inferred at 1.5x coverage compared to higher coverage for clones where it exists.

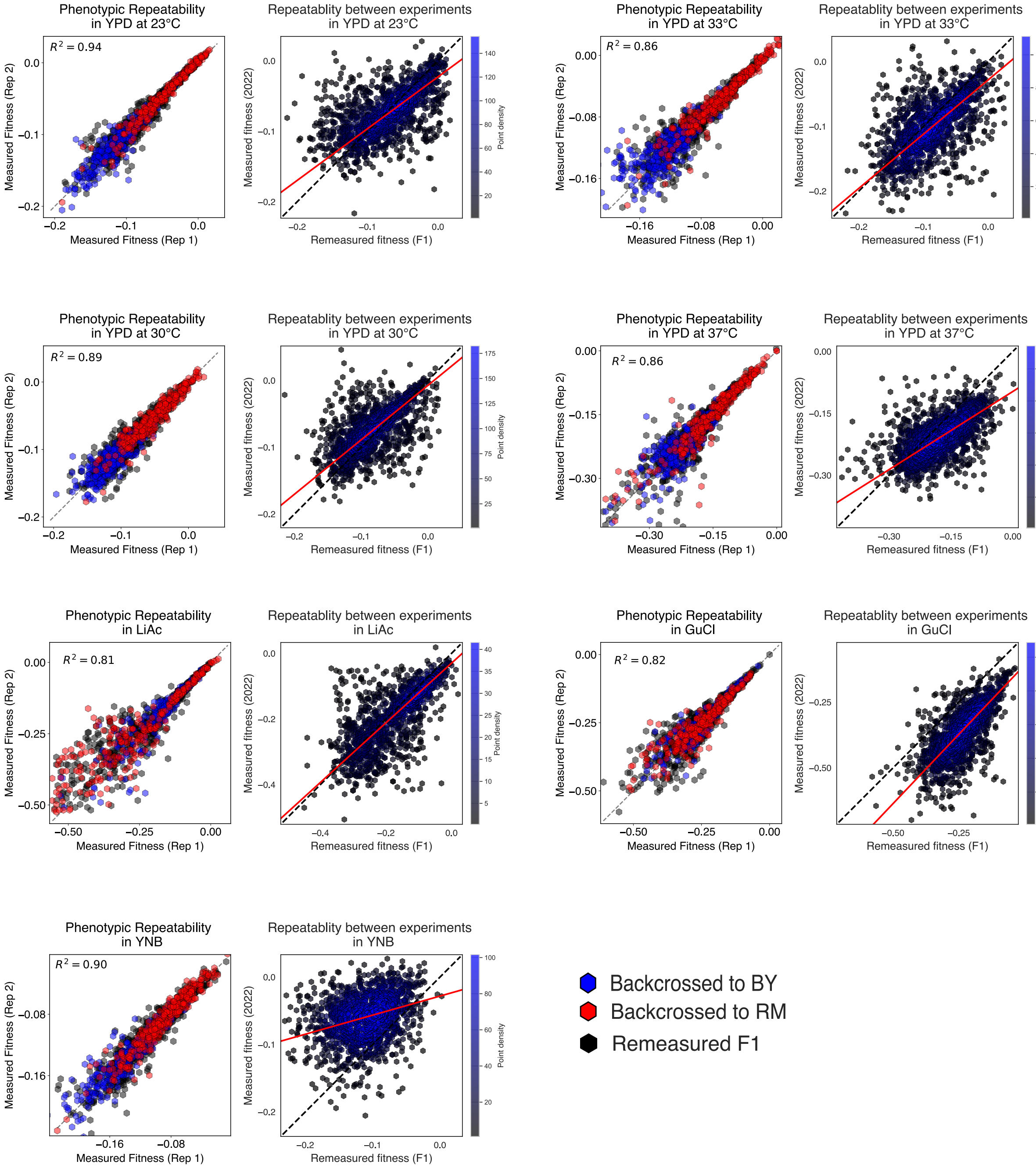


**Fig. S2.** Phenotypic measurements appear robust, both within and between experiments. Each growth environment was measured in 2 replicates. When compared to each other, we get good inter-replicate correlations, with $R^2 > 0.8$ in all cases. For the main analysis, fitness was inferred jointly with both replicates. The best fit line is plotted weighted by the density of points. Since the F1 segregants from our panel were also present in the original 2022 study, we can also directly compare the remeasured fitness values. The 2 different measurements are in agreement with each other. The red line is the best fit line to all data, weighted by the density of points (higher density represented by blue). YNB is a minimal growth media, often stressful for the yeast. The distribution of fitness values is also much smaller in this environment. Fitness inference is noisier in this environment, potentially explaining the different between experiments. Our fitness inference procedure can also be sensitive to population composition of all strains, explaining the slight deviations seen.

**SI Dataset S1 (primers_and_sequences.csv)**
Details of primers used during construction and custom sequencing pipelines

**SI Dataset S2 (segregant_info_coverage.csv)**
Segregant IDs along with raw sequencing coverage

### References


1. AN Nguyen Ba, et al., Barcoded bulk qtl mapping reveals highly polygenic and epistatic architecture of complex traits in yeast. *eLife* **11**, e73983 (2022).
2. RB Brem, G Yvert, R Clinton, L Kruglyak, Genetic dissection of transcriptional regulation in *saccharomyces cerevisiae*. *Science* **296**, 752–755 (2002).
3. AN Nguyen Ba, et al., High-resolution lineage tracking reveals travelling wave of adaptation in laboratory yeast. *Nature* **575**, 494–499 (2019) Epub 2019 Nov 13.
4. RD Gietz, RH Schiestl, Quick and easy yeast transformation using the liac/ss carrier dna/peg method. *Nat. Protoc.* **2**, 35–37 (2007).
5. MS Johnson, et al., Phenotypic and molecular evolution across 10,000 generations in laboratory budding yeast populations. *eLife* **10**, e63910 (2021).
6. M Baym, et al., Inexpensive multiplexed library preparation for megabase-sized genomes. *PLOS ONE* **10**, e0128036 (2015).
7. KW Broman, H Wu, Ś Sen, GA Churchill, R/qtl: Qtl mapping in experimental crosses. *bioinformatics* **19**, 889–890 (2003).
8. B Langmead, S Salzberg, Fast gapped-read alignment with bowtie 2. nat 936 methods 9, 357–359 (2012).
9. AM Smith, et al., Quantitative phenotyping via deep barcode sequencing. *Genome Res.* **19**, 1836–1842 (2009).
10. RA Fisher, The correlation between relatives on the supposition of mendelian inheritance. *Transactions Royal Soc. Edinb.* **52**, 399–433 (1918).
11. S Wright, Coefficients of inbreeding and relationship. *The Am. Nat.* **56**, 330–338 (1922).
